# Effects of bimodal divided attention on cortical representations of linguistic context during continuous speech perception in noise

**DOI:** 10.64898/2026.04.28.721419

**Authors:** Zilong Xie

## Abstract

Speech perception often takes place in environments with competing sensory inputs, both within the auditory modality and across modalities; for example, following a conversation in a noisy cafe while simultaneously reading a menu. This study examined the extent to which dividing attention between auditory and visual modalities (bimodal divided attention) influences linguistic context processing across hierarchical levels during continuous speech perception in noise. Electroencephalographic (EEG) responses were recorded while participants listened to audiobook stories in multitalker babble as a secondary task, concurrently performing a demanding primary visual task that imposed either low or high cognitive load. Behaviorally, speech comprehension accuracy was significantly lower under high-load than low-load dual-task conditions. Multivariate temporal response function (mTRF) encoding models were used to predict EEG responses from information-theoretic measures (entropy and surprisal) indexing linguistic context at sublexical, word-form, and sentence levels. Significant neutral tracking was observed at the word-form and sentence levels, but not the sublexical level. Critically, neutral tracking of sentence-level linguistic representations was significantly reduced under high compared to low load, with effects emerging at latencies beyond 200 ms. In contrast, neutral tracking of word-form-level representations was unaffected by dual-task load. mTRF analyses further revealed that neutral tracking of acoustic features was not modulated by dual-task load. These findings indicate that bimodal divided attention selectively disrupts cortical representations of sentence-level linguistic context, while lower-level processing remains relatively preserved. Such impairments in higher-level linguistic processing may contribute to reduced speech comprehension during multitasking in noisy environments.

## Introduction

In everyday life, speech perception often unfolds amid competing sensory inputs, both within the auditory modality and across other modalities; for example, following a conversation in a noisy cafe while simultaneously reading a menu. Such complex listening environments place substantial demands on perceptual and cognitive resources and can lead to communication breakdowns. Successful speech perception under these conditions requires listeners to allocate attention to the auditory modality while managing demands from nonauditory inputs, to segregate target speech from background noise, and to engage selective attention mechanisms that enhance target speech while suppressing interfering sounds (Shinn-Cunningham, 2008). The present study investigated the neural mechanisms underlying the effects of dividing attention between modalities—specifically audition and vision (i.e., bimodal divided attention)—on the processing of natural continuous speech in noise.

Speech processing entails the extraction and encoding of acoustic information from the speech signal and its progressive transformation into linguistic representations of increasing complexity (Brodbeck & Simon, 2020; Hickok & Poeppel, 2007). A key mechanism proposed to support this transformation is the use of linguistic context across multiple hierarchical levels to facilitate the interpretation of incoming speech input (Kuperberg & Jaeger, 2016; Ryskin & Nieuwland, 2023). Research on linguistic context processing has traditionally relied on paradigms employing highly controlled, constructed stimuli, such as isolated words (e.g., Ganong, 1980; Giovannone & Theodore, 2021; Gwilliams et al., 2018; Lam et al., 2017; Sohoglu et al., 2024) or sentences (e.g., Connine et al., 1991; Hsin et al., 2023; Hsin & Lee, 2024; Obleser & Kotz, 2011; Sarrett et al., 2020; Strauß et al., 2013; Xie, 2025). Over the past decade, however, there has been a growing shift toward paradigms that use natural, continuous speech, which more closely approximates real-word speech and language processing (Brodbeck & Simon, 2020; Gillis et al., 2022; Hamilton & Huth, 2020).

Of relevance to the present study, Brodbeck et al. (2022) asked participants to listen to continuous narrative speech (i.e., audiobook stories) under quiet listening conditions while neural responses were recorded. They examined how linguistic context at three hierarchical levels modulated neural responses to phonemes. The sublexical-level context incorporated local phoneme context (i.e., the preceding four phonemes) to predict upcoming phonemes. The word-form-level context, grounded in the cohort model of word recognition (Marslen-Wilson, 1987), incorporated word statistics (i.e., the probability of words with a given phoneme sequence in the lexicon) to predict upcoming phonemes. The sentence-level context extended the word-form model by additionally incorporating sentence context (i.e., the four preceding words) to estimate word-level statistics. The authors found that linguistic context at all three levels significantly modulated neural responses to phonemes. Effects for all context levels were observed in the superior temporal gyrus, with the sentence-level context additionally engaging more ventral regions of the temporal lobe. In terms of temporal dynamics, all context levels modulated early neural responses within the first 100 ms; however, only the sentence-level context influenced later responses, emerging after approximately 150 ms and persisting up to ∼500 ms. Together, these findings suggest that continuous speech processing involves the parallel use of linguistic context across multiple hierarchical levels, including sublexical-, word-form-, and sentence-level representations (see also, e.g., Brodbeck et al., 2024; Karunathilake et al., 2025).

However, because this and other related work have focused on speech processing under auditory-only attention, it remains unclear whether and how bimodal divided attention modulates linguistic context processing across these hierarchical levels. Addressing this issue is essential for advancing our understanding of the role of domain-general cognitive processes in linguistic prediction during speech and language comprehension (Ryskin et al., 2020; Ryskin & Nieuwland, 2023). Recently, Xie et al. (2023) examined this question using electroencephalography (EEG) in conjunction with an audiovisual dual-task paradigm. Bimodal divided attention was operationalized by requiring participants to perform a demanding primary visual task under either low or high cognitive load while concurrently listening to continuous speech presented in quiet as a secondary task. These two dual-task conditions were compared with an auditory single-task condition, in which participants focused exclusively on the speech stimuli while disregarding concurrent visual stimuli.

Xie et al. (2023) replicated the core findings from Brodbeck et al. (2022), demonstrating that linguistic context at sublexical, word-form, and sentence levels significantly modulated neural responses to phonemes. Crucially, neural responses reflecting these context effects were stronger in the auditory single-task condition than in either dual-task condition, with no significant difference observed between the low- and high-load dual-task conditions. Although divided attention across modalities attenuated context-related neural responses, these effects were not eliminated under dual-task conditions. Moreover, the magnitude and pattern of attention-related modulation was comparable across all three levels of linguistic context. Collectively, these findings suggest that bimodal divided attention may reduce—but does not abolish—the use of linguistic context during continuous speech processing and that its influence appears relatively uniform across hierarchical levels of linguistic representations.

Notably, however, this study examined speech processing exclusively under quiet listening conditions. As a result, it remains unclear to what extent additional processing demands imposed by background noise further influence the attentional modulation of linguistic context processing in crossmodal settings. Prior work examining attentional effects on linguistic context processing during continuous speech perception in noise have largely emphasized auditory selective attention and has rarely explicitly tested attentional effects across multiple hierarchical levels of linguistic context processing. For example, several studies have used a two-talker paradigm in which listeners selectively attended to one of two competing talkers narrating different audiobook stories. One study focused on neural responses indexing sentence-level context processing and showed that these responses were significantly stronger for attended speech than for unattended speech (Broderick et al., 2018). Another study examined neural responses indexing sublexical-level context processing and demonstrated that these responses were robust for attended speech but were abolished for unattended speech (Brodbeck et al., 2018a). Importantly, selective and divided attention are thought to be supported by distinct underlying mechanisms (Johnson & Zatorre, 2006; Loose et al., 2003), and their effects on continuous speech processing may therefore differ (e.g., selective attention in Brodbeck et al., 2018 versus divided attention in Xie et al., 2023). Accordingly, the present study aims to investigate how bimodal divided attention modulates linguistic context processing across multiple hierarchical levels during continuous speech perception in noise.

Here, we adopted the audiovisual dual-task paradigm from Xie et al. (2023) to manipulate bimodal divided attention. Participants performed a demanding primary visual task under either low or high cognitive load while concurrently listening to continuous speech in noise as a secondary task. We compared the two dual-task conditions (low versus high load) to assess the effects of bimodal divided attention on linguistic context processing. We tested the hypothesis that sentence-level context processing would be more susceptible to attentional modulation than word-form and sublexical context processing, owing to its reliance on tracking and integrating contextual information over longer timescales, which may necessitate the involvement of cognitive resources (Alexander & Brown, 2018; Miller & Cohen, 2001; Shain et al., 2022).

## Materials & Methods

### Participants

The study included 24 younger adults (20 females) aged 18-23, recruited from the Tallahassee, Florida, area. The sample size was determined based on previous research on bimodal attention and its effects on the neural processing of speech (e.g., Gennari et al., 2018; Kasper et al., 2014; Xie et al., 2023). All participants had normal hearing and normal or corrected-to-normal vision. Normal hearing was defined as having pure-tone air-conduction thresholds ≤25 dB hearing level at octave frequencies from 0.25 to 8 kHz in both ears, assessed using an Interacoustics Equinox 2.0 PC-based audiometer (Interacoustics, Middelfart, Denmark). All participants were native speakers of English; four reported speaking a second language (three in Spanish and one in Hindi). No participant reported a history of head trauma. The study protocol was approved by the Institutional Review Board at Florida State University (IRB#: STUDY00003674). Written informed consent was obtained from all participants, who received monetary compensation for their participation.

### Experimental design and stimuli

The experimental design and stimuli were adapted from Xie et al. (2023); readers are referred to that work for additional details.

In this study, participants completed a dual-task paradigm comprising a primary visuospatial n-back task of varying cognitive load and a secondary task of listening to continuous speech in noise. Visual stimuli were blue squares appearing at one of eight positions surrounding a central fixation cross on a black background. Auditory target stimuli were approximately 60-second excerpts from the audiobook *Alice’s Adventures in Wonderland* (http://librivox.org/alices-adventures-in-wonderland-by-lewis-carroll-5), narrated by an adult male speaker of American English at a 22.05 kHz sampling rate. Each excerpt was resampled to 20 kHz and calibrated to 65 dB sound pressure level. The background noise was multi-talker babble originally described by Varga and Steeneken (1993) and obtained from Bechtold (2020). This recording features 100 people conversing in a canteen, sampled at 19,980 Hz with a total duration of approximately 235 seconds. For this study, the noise was resampled to match the speech stimuli, and the first 69 seconds were extracted. A 10-ms ramp was applied to the onset and offset. This segment was then mixed with the story excerpts at a signal-to-noise ratio of 5 dB. To help participants focus on the target speech, it began 3 seconds after the onset of the noise and ended before the noise segment concluded.

Visual stimuli were displayed on a 23.8-inch monitor (resolution: 1920 × 1080, refresh rate: 60 Hz) positioned approximately 150 cm from participants. Auditory stimuli were presented monaurally to the right ear through ER-2 insert earphones (Etymotic Research, Elk Grove Village, IL). Monaural presentation was chosen because this study was part of a larger investigation on aging and continuous speech processing, and this approach increases the likelihood of recruiting older adults with clinically normal hearing.

Cognitive load was manipulated using 3- and 0-back visual tasks. In the high-load condition (3-back), participants responded when the current square matched the one presented three positions back earlier. In the low-load condition (0-back), they responded when the square matched the first in the sequence. The auditory task remained constant across conditions: participants answered two multiple-choice comprehension questions about the story segment at the end of each trial.

Each condition comprised 15 trials, pairing unique visual sequences with distinct story segments. The two conditions were presented in separate blocks, with block order counterbalanced across participants. At the start of each condition, two practice trials were administered. These practice trials were similar to the actual task, except that the auditory stimuli consisted of story segments not used in the actual trials. The same auditory stimuli were used for practice in both the low- and high-load conditions. The experiment was programmed and controlled using E-Prime 3.0.3.214 (Schneider et al., 2002).

At the end of each condition, participants rated the effort required to complete the task on a 10-point scale (1 = no effort, 10 = extreme effort) and estimated the percentage of the story they understood (0–100%). The high-load condition was associated with significantly greater perceived effort and lower story understanding compared to the low-load condition: Effort: high-load (mean = 8.5, SD = 1.50) vs. low-load (mean = 4.83, SD = 2.22), t(23) = 10.498, *p* < 0.001; Story understanding: high-load (mean = 49.6%, SD = 23.4%) vs. low-load (mean = 77.0%, SD = 18.0%), t(23) = −5.983, *p* < 0.001.

Following the experiment, participants completed an exit questionnaire to assess their familiarity with the story segments. They indicated whether they had previously read *Alice’s Adventures in Wonderland* or watched a film adaptation, and if so, approximately when. They also rated the extent to which prior exposure aided task performance on a 10-point scale (1 = not at all, 10 = very much). Of the 24 participants, nine reported having read the book, and sixteen had seen the movie. A two-way mixed ANOVA examined whether prior familiarity (book or movie) influenced auditory task accuracy under low- and high-load conditions. No significant main effects of reading (*p* = 0.899, η²_p_ = 0.0006) or watching (*p* = 0.47, η²_p_ = 0.019) were observed, nor were any interactions with task condition significant (*p*s > 0.72, η²_p_ < 0.002), indicating that prior familiarity did not meaningfully affect performance.

### Electrophysiological data acquisition and preprocessing

#### Acquisition

The experiment took place in a dimly lit, sound-attenuating booth. Participants were seated comfortably throughout the tasks. Electroencephalography (EEG) signals were recorded at a sampling rate of 5 kHz using an Easycap recording cap (www.easycap.de) equipped with 64 actiCAP active electrodes (Brain Products, Gilching, Munich, Germany). Electrode placement followed the extended 10-20 system (Oostenveld & Praamstra, 2001), with the Fpz site serving as the ground and Fz as the reference. All electrode impedances were maintained below 25 kΩ. EEG data were acquired using a Brainvision actiCHamp Plus amplifier linked to BrainVision Recorder software V. 1.25.0001.

#### Preprocessing

The raw EEG data were preprocessed offline using custom scripts implemented in MNE-Python (Gramfort et al., 2013) and EEGLAB (Delorme & Makeig, 2004). The raw data were re-referenced to the average of electrodes TP9 and TP10, followed by band-pass filtering between 1 and 15 Hz using a Hamming-windowed sinc FIR filter with default EEGLAB settings. The data were then segmented into 66-second epochs, time-locked to the onset of the corresponding story segments, and downsampled to 500 Hz. Independent component analysis was applied to the epoched data pooled across both task conditions for each participant using the Infomax algorithm. Artifact-related components, specifically those associated with muscle activity and eye movements, were automatically detected using ICLabel (Pion-Tonachini et al., 2019) with threshold values between 0.8 and 1.0 and subsequently removed.

#### Assessing neural tracking of visual and auditory stimuli

The multivariate temporal response function (mTRF) approach (Crosse et al., 2016; Ding & Simon, 2012), implemented via the Eelbrain Python toolkit (Brodbeck et al., 2023) was used to assess neural tracking of visual and auditory stimuli. This technique employs time-delayed multiple regression to predict EEG signals from stimulus-derived predictor variables. The predictive power of each predictor set (e.g., visual features) was interpreted as an index of the strength of the corresponding neural representation. Examples of these predictors are shown in Figure 1. Visual and acoustic predictors were estimated following procedures similar to those described in Xie et al. (2023), while linguistic predictors were directly adopted from that work. A brief overview of these predictors is provided below; for additional methodological details, see Xie et al. (2023). To enhance computational efficiency, both EEG data and predictor variables were downsampled to 100 Hz before the mTRF analysis. Additionally, only the first 56 seconds of each trial were included for the mTRF analysis to match the shortest trial duration.

**Figure 1.**
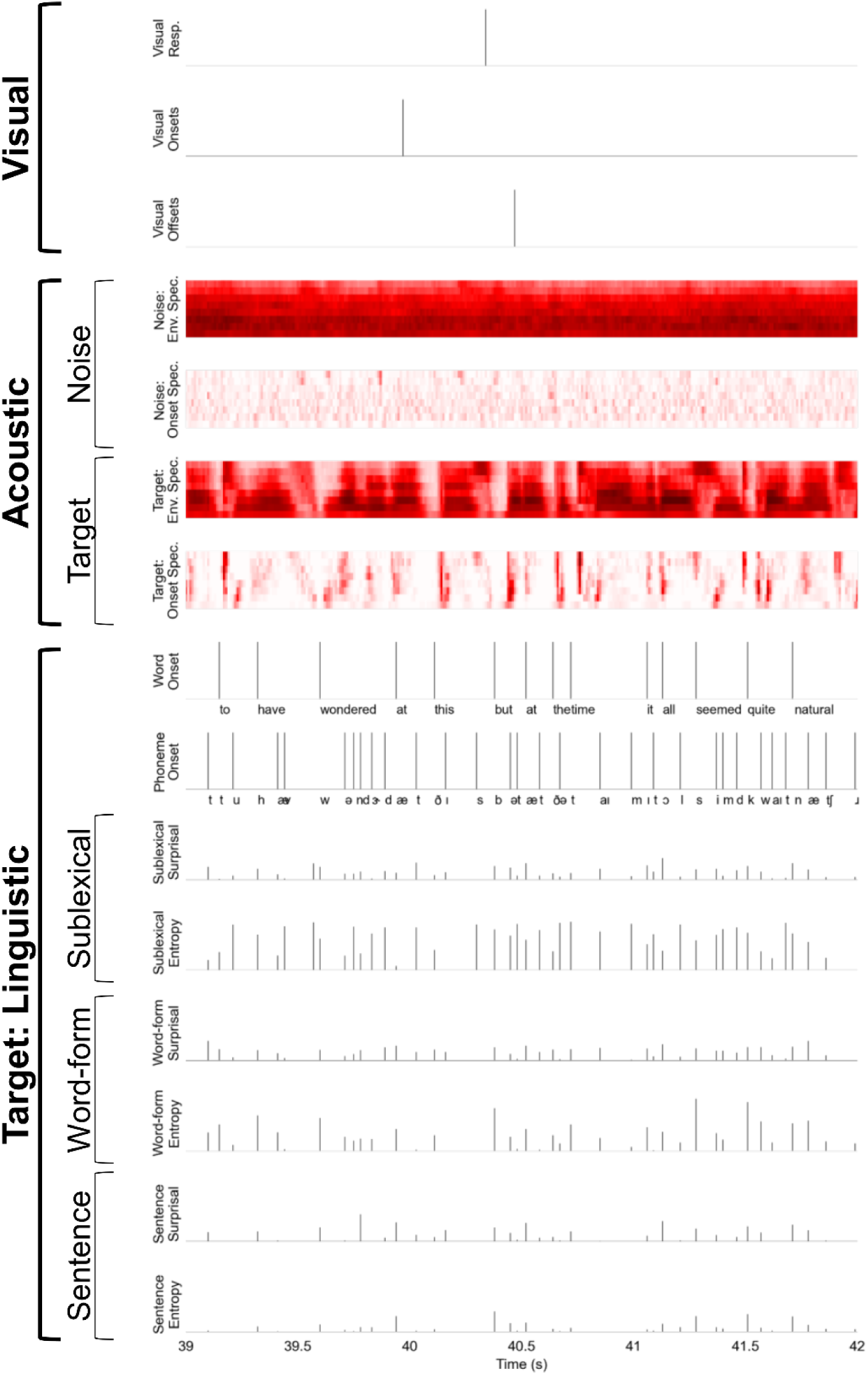
Examples of visual, acoustic, and linguistic predictors used to model the EEG responses in the mTRF analysis.

### Visual and acoustic predictors

The visual predictor consisted of a one-dimensional time series with an impulse at the onset and offset of each blue square. To account for potential neural activity related to participants’ responses, an additional one-dimensional time series was included, containing impulses at time points when participants responded to the visual stimuli.

Acoustic predictors were designed to capture the spectro-temporal properties of both the target speech and background noise. These predictors were derived from high-resolution gammatone spectrograms spanning 256 frequency bands from 20 to 10,000 Hz. An *envelope spectrogram* predictor was obtained by summing the 256-band spectrogram into eight logarithmically spaced frequency bands. Additionally, an *onset spectrogram* predictor was created by applying an auditory edge detection model (Brodbeck et al., 2020; Fishbach et al., 2001) to each band of the 256-band spectrograms, and then summing them into eight logarithmically spaced frequency bands.

### Linguistic predictors

Linguistic predictors were computed only for the target speech, comprising information-theoretic measures—surprisal and entropy—of phoneme sequences using sublexical, word-form, and sentence models (briefly described below). These measures were expressed as time series with impulses of varying magnitude at each phoneme onset. Phoneme sequences and their onsets were obtained from the speech stimuli through forced alignment with the Montreal Forced Aligner (McAuliffe et al., 2017).

#### Sublexical model

The sublexical model uses a 5-gram model (Heafield, 2011) trained on phoneme sequences from all sentences in the SUBTLEX-US corpus (Keuleers et al., 2010), without word boundary removed. It predicts each current phoneme from the four preceding ones and was applied to target phoneme sequences to generate *sublexical surprisal* and *entropy* predictors. *Sublexical surprisal* represents the level of probabilistic surprisal of the current phoneme given its four-phoneme context, while *sublexical entropy* captures the uncertainty about the next phoneme based on the same context. These predictors are limited to simple phoneme sequence statistics and do not account for word boundaries (Brodbeck et al., 2022; Vitevitch & Luce, 1999, 2016).

#### Word-form model

The word-form model is grounded in the cohort model of word recognition (Brodbeck et al., 2018b; Marslen-Wilson, 1987). Each word was assigned a prior probability based on its count frequency in the SUBTLEX corpus (Keuleers et al., 2010), with a default value of 1 for words not present in the corpus. For each word in the target speech, the model begins with the full lexicon and progressively removes words that do not match the observed phoneme sequence as each phoneme is processed. The evolving probability distribution over the lexicon at each phoneme position was used to compute *word-form surprisal* and *entropy* predictors. *Word-form surprisal* reflects the level of probabilistic surprisal of the current phoneme given the preceding phonemes within the word, while *word-form entropy* represents the uncertainty of all words currently in the cohort that match the observed phoneme sequence. The lexicon was constructed by combining pronunciations from the Montreal Forced Aligner English dictionary and the Carnegie Mellon University Pronouncing Dictionary (http://www.speech.cs.cmu.edu/cgi-bin/cmudict).

#### Sentence model

The sentence model applies the same procedure to the word-form model to compute *sentence surprisal* and *entropy* predictors, with one key difference: the prior probability of each word was defined from the four preceding words, using a lexical 5-gram model (Heafield, 2011) trained on the whole SUBTLEX-US corpus (Keuleers et al., 2010).

Thus, sentence-level predictors incorporate a broader linguistic context, spanning multiple words, compared to sublexical and word-form predictors. Further, to account for the effects of linguistic segmentation, two additional predictors were included: a word-onset predictor, assigning a unit impulse at the onset of each word-initial phoneme, and a phoneme-onset predictor, assigning a unit impulse at the onsets of all other phonemes.

### Estimation of neural tracking

This study examined to extent to which neutral tracking of visual information and the acoustic and linguistic properties of target speech differ across the two task conditions. For each participant and condition, a forward encoding mTRF model was applied, using the previously defined predictors to model EEG responses at individual electrodes. Models were estimated using a boosting algorithm with 5-fold cross-validation, implemented in Eelbrain. The mTRFs were parameterized with 50-ms Hamming basis functions spanning the stimulus-EEG lag from −100 to 500 ms, and all predictors were jointly optimized via coordinate descent to minimize ℓ_2_ loss.

To implement 5-fold cross-validation, the data (n = 15 trials) were partitioned into 5 folds (3 trials per fold). In each iteration, one fold was held out for testing, while the remaining folds rotated as the validation set, yielding four mTRFs per testing fold. These mTRFs were averaged to produce a single mTRF for that iteration, which was then used to predict EEG in the held-out (testing) fold. Predicted EEG responses from all five testing folds were concatenated to estimate a per-electrode measure of model performance, i.e., the proportion of the variation in the actual EEG response that is explained by the model (i.e., % explained variance). Final mTRFs per electrode were obtained by averaging the mTRFs across all testing folds.

To quantify neural tracking of specific speech features, we contrasted a full model with reduced models in which the predictor(s) of interest were removed. The full model included visual predictors (onsets, offsets, and responses), acoustic predictors for background noise (envelope and onset spectrograms), acoustic predictors for target speech (envelope and onset spectrograms), and linguistic predictors (sublexical surprisal and entropy; word-form surprisal and entropy; sentence surprisal and entropy; plus word and phoneme onsets). Visual tracking was assessed by removing visual onsets and offsets; acoustic tracking of the target speech by removing the target envelope and onset spectrograms; sublexical-level processing by removing sublexical surprisal and entropy; word form-level processing by removing word form surprisal and entropy; and sentence level processing by removing sentence surprisal and entropy. The change in % explained variance (i.e., Δ% explained) between the full and reduced models was used to index the strength of neutral tracking for the corresponding predictor(s).

#### Statistical analysis

All statistical analyses were performed in Eelbrain, unless otherwise noted.

First, we examined the impact of task condition (low vs. high load) on behavioral performance and on neural tracking of visual, acoustic, and linguistic properties, using one-sided paired t-tests with an alpha level of .05. Effect sizes (Cohen’s *d*) and Bayes Factors (BF) were also estimated. Additional details are provided below.

Behavioral performance was indexed by three measures: visual accuracy, visual reaction time (RT), and auditory accuracy. Visual accuracy was calculated as the hit rate (i.e., correct responses to targets) minus the false alarm rate (i.e., incorrect identification of non-targets as targets). Visual RT was defined as the median RT for hits only. Auditory accuracy was expressed as the percentage of correctly answered story comprehension questions. The analyses were performed in R (version 4.5.1; Team, 2022), and BF values were estimated using the BayesFactor package (version 0.9.12.4.7; Morey et al., 2022).

Neural tracking was quantified by the Δ% explained by each predictor or predictor set. The Δ% explained values were averaged across task conditions and tested against zero using a one-sided, cluster-based permutation test (Maris & Oostenveld, 2007). Clusters were identified using a t-threshold corresponding to uncorrected p ≤ 0.05, and corrected p-values were derived from a null distribution of 10,000 permutations (or all possible permutations if fewer). The largest t-value within each cluster (tmax) was reported as an effect size estimate (Brodbeck et al., 2018a). Significant clusters were treated as regions of interest (ROIs), and Δ% explained values were averaged within these ROIs for subsequent condition comparisons. Further, when a significant task-condition effect was observed, follow-up tests tested whether neural tracking for the relevant predictor(s) persisted within each condition, by testing Δ% explained values against zero using one-sided one-sample t-tests.

Finally, we analyzed the effect of task conditions on TRFs for visual, acoustic, and linguistic predictors. The TRFs were extracted from the full model, and the global field power (GFP) was computed across the corresponding ROI(s) identified above. GFP values were compared between conditions using two-sided, cluster-based permutation tests in Eelbrain with default parameters, except for the analysis time window. For visual predictors, TRFs for visual onsets and offsets were concatenated to capture responses to visual stimuli as a whole, resulting in TRFs spanning with lags from −100 to 1000 ms relative to stimulus onset because the visual stimuli lasted exactly 500 ms. The analysis window was 0–000 ms for visual predictors and 0–500 ms for acoustic and linguistic predictors. For the acoustic predictors, TRFs were averaged across the eight frequency bands before analysis.

## Results

Table 1 summarizes the key findings regarding the effects of task conditions on behavioral performance and neural visual, acoustic, and linguistic processing.

**Table 1.**
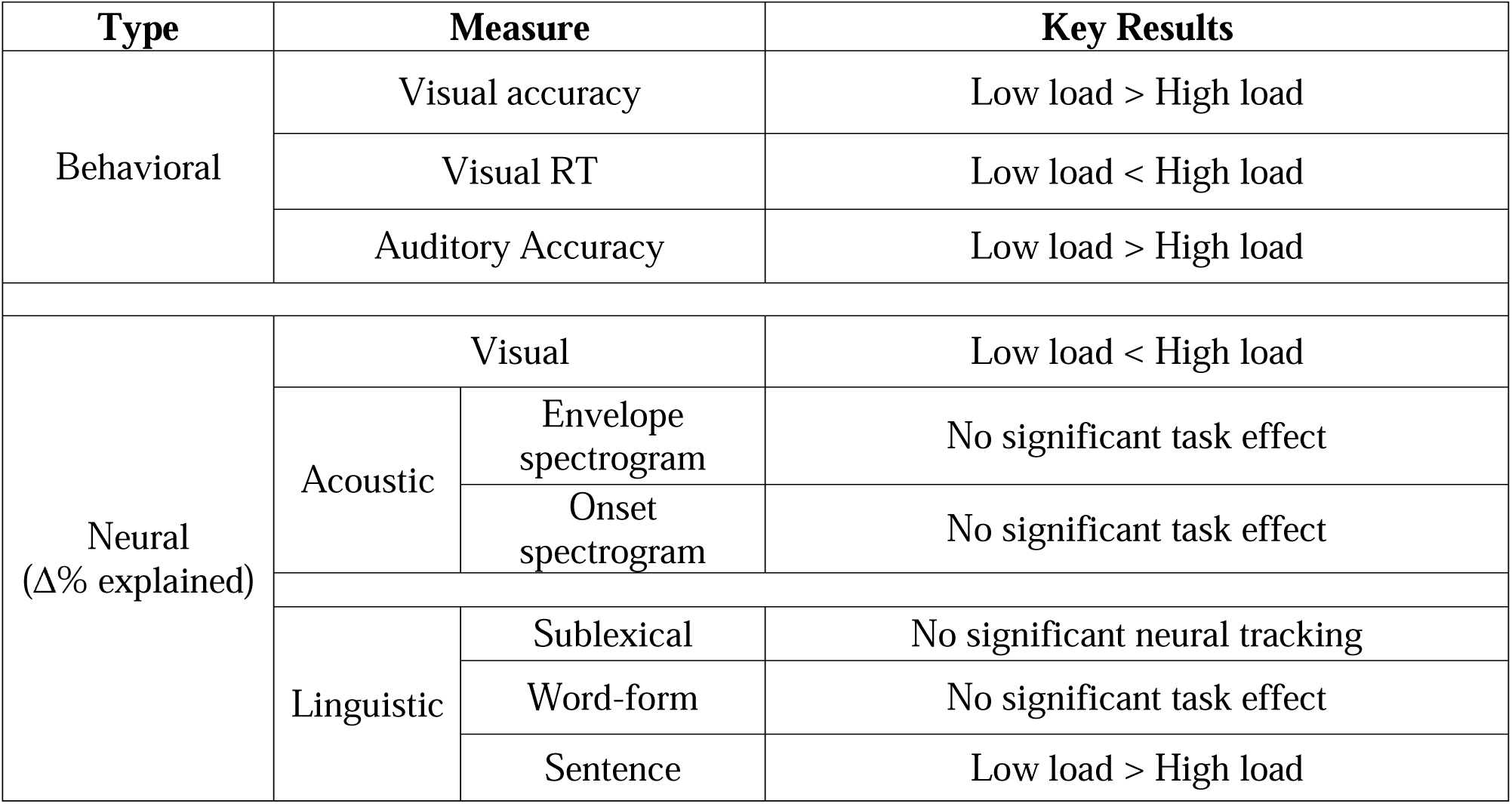
Task Effects on Continuous Speech Processing.

| Type | Measure |  | Key Results |
| --- | --- | --- | --- |
| Behavioral | Visual accuracy |  | Low load > High load |
|  | Visual RT |  | Low load < High load |
|  | Auditory Accuracy |  | Low load > High load |
| Neural<br>(Δ% explained) | Visual |  | Low load < High load |
|  | Acoustic | Envelope spectrogram | No significant task effect |
|  |  | Onset spectrogram | No significant task effect |
|  | Linguistic | Sublexical | No significant neural tracking |
|  |  | Word-form | No significant task effect |
|  |  | Sentence | Low load > High load |

### Visual task load impairs behavioral visual and auditory performance

Figure 2A and 2B display the accuracy and RT of the visual task for individual participants. Compared to the low load (0-back) condition, the high load (3-back) condition was associated with lower accuracy [*t* (23) = 8.791, *p* < 0.001, Cohen’s d = 1.79, BF = 1.55×10^6^] and slower RT [*t* (23) = −6.866, *p* < 0.001, Cohen’s d = 1.40, BF = 3.21×10^4^]. These results confirmed that the manipulation of cognitive load in the visual task was successful.

**Figure 2.**
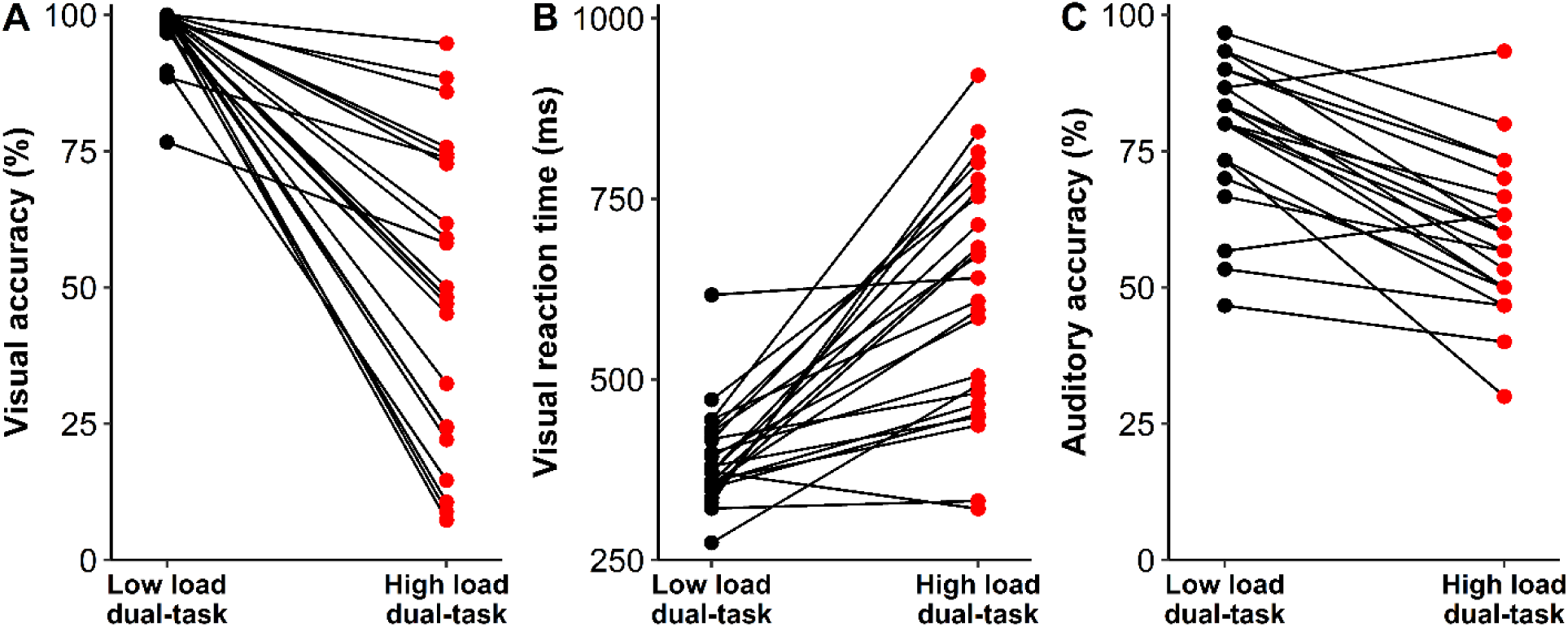
Behavioral performance on visual and auditory tasks. (A) Accuracy for the low load (0-back) and high load (3-back) visual tasks, computed as hit rate (i.e., correct target responses) minus false alarm rate (i.e., identifying a non-target as being a target). (B) Median reaction time (RTs) for hits only (correct target responses) in the low- and high-load visual tasks. (C) Accuracy on the auditory task, quantified as the proportion of story comprehension questions answered correctly. Each line in panels (A to C) corresponds to an individual participant (n = 24).

Figure 2C displays the auditory task accuracy for individual participants. The mean accuracy was 79.4% (*SD* = 13.0%) in the low load dual-task condition and 60.2% (*SD* = 13.7%) in the high load dual-task condition. The effect of task condition was significant [*t* (23) = 7.981, *p* < 0.001, Cohen’s d = 1.63, BF = 3.18×10^5^].

Further, we examined the relationship between visual and auditory task performance during the dual-task conditions. The change in auditory accuracy [i.e., (low load – high load)/low load] was not significantly correlated with the change in visual RT [i.e., (high load – low load)/low load] (Spearman’s ρ = −0.08, *p* = 0.707, BF = 0.448) or the change in visual accuracy (Spearman’s ρ = 0.074, *p* = 0.733, BF = 0.461).

### Neural tracking of sentence-level linguistic information reduced with visual task load, but not at sublexical or word-form levels

The sentence-level linguistic predictors significantly enhanced model performance beyond visual, acoustic, and other linguistic predictors, with a small cluster at central regions (*t*_max_ = 3.94, *p* = .008; Figure 3A). These results provide evidence for robust cortical tracking of sentence-level linguistic information. Furthermore, as illustrated in Figure 3B, the high load dual-task condition yielded significantly lower predictive power (mean = 0.006, *SD* = 0.013) than the low load dual-task condition (mean = 0.012, *SD* = 0.012) [*t*(23) = 1.818, *p* = 0.041, Cohen’s d = 0.499, BF = 1.771]. These results suggest that increased visual task load impairs neural tracking of linguistic information at the sentence-level. Finally, to understand how differences in model predictive power manifest in the cortical responses, we compared the mTRFs between the two dual-task conditions. In line with the findings on predictive power, as shown in Figure 3C, the high load dual-task condition produced significantly smaller mTRF amplitudes than the low load dual-task condition from 220 to 300 ms (*p* = 0.005).

**Figure 3.**
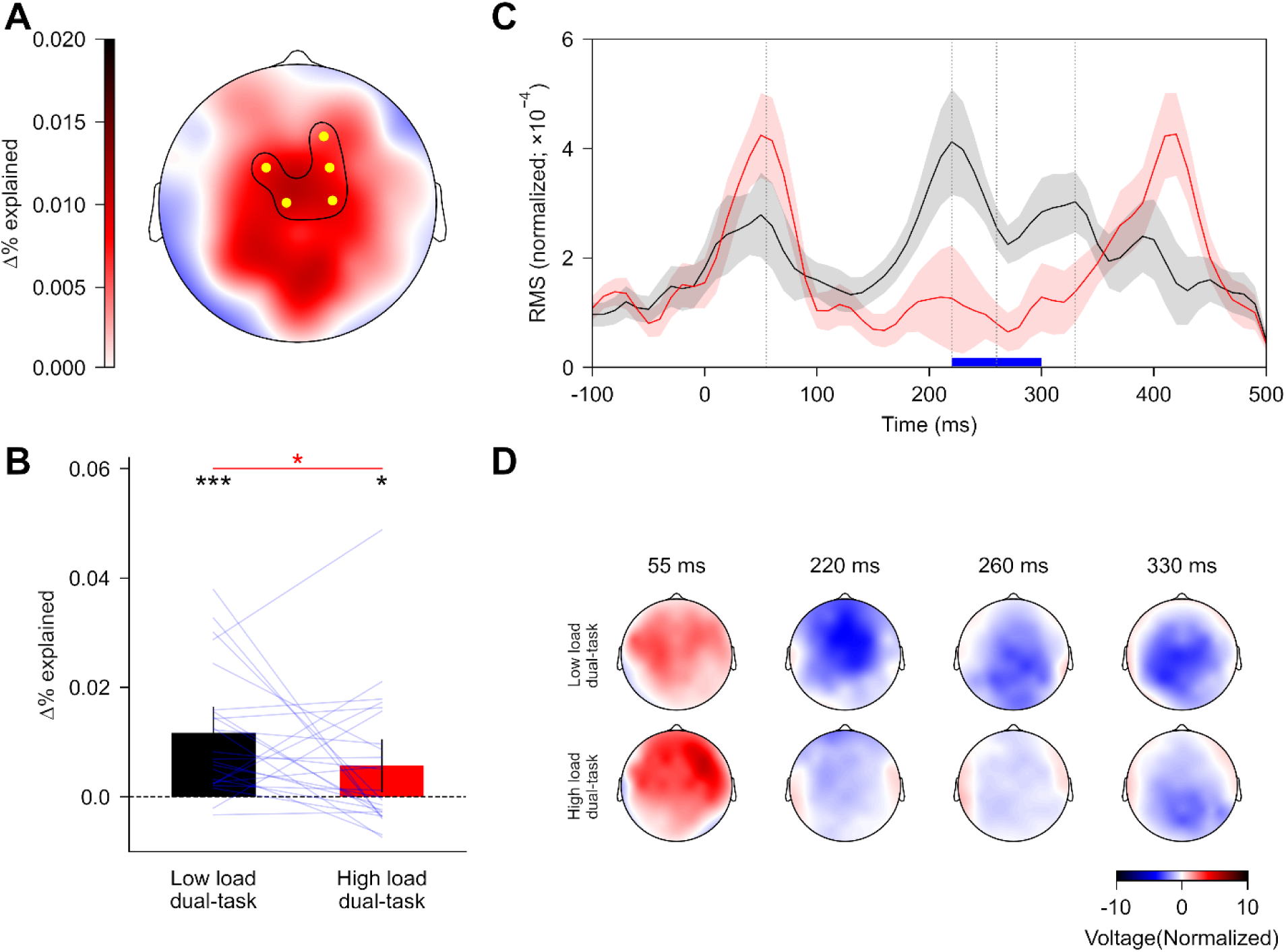
Neural tracking of sentence-level linguistic information across dual-task conditions. (A) The topographic map illustrates the increase in prediction power (expressed as Δ% variance explained) attributed to sentence-level linguistic predictors (sentence surprisal and entropy). A statistically significant increase was observed within a single cluster, marked by the highlighted yellow electrodes. (B) Prediction power across dual-task conditions, with blue lines representing individual participants. A red asterisk indicates the statistical significance of the difference between conditions, while a black asterisk marks whether the prediction power for each condition was significantly above zero. Error bars reflect the 95% within-subject confidence interval. (C) The global field power of the TRFs for the sentence-level predictors. The TRFs were derived by averaging responses across the surprisal and entropy predictors. Shaded regions represent within-subject standard errors around the mean (color coding as in panel B). A blue horizontal line highlights the time interval where TRFs significantly differed between conditions. (D) Scalp topographies at selected time points (indicated by grey vertical lines in panel C). * *p* < 0.05, *** *p* < 0.001.

The word-form level predictors significantly enhanced model performance beyond visual, acoustic, and other linguistic predictors, with a cluster at central regions (*t*_max_ = 5.09, *p* < 0.001; Figure 4A). These results provide evidence for robust neural tracking of linguistic information at the word-form level. Additionally, task condition did not significantly influence the predictive power for the word-form level predictors [*t*(23) = 0.618, *p* = 0.271, Cohen’s d = 0.176, BF = 0.511; Figure 4B]. Finally, the mTRFs for the word-form level predictors (Figure 4C) were not significantly different between the two dual-task conditions.

**Figure 4.**
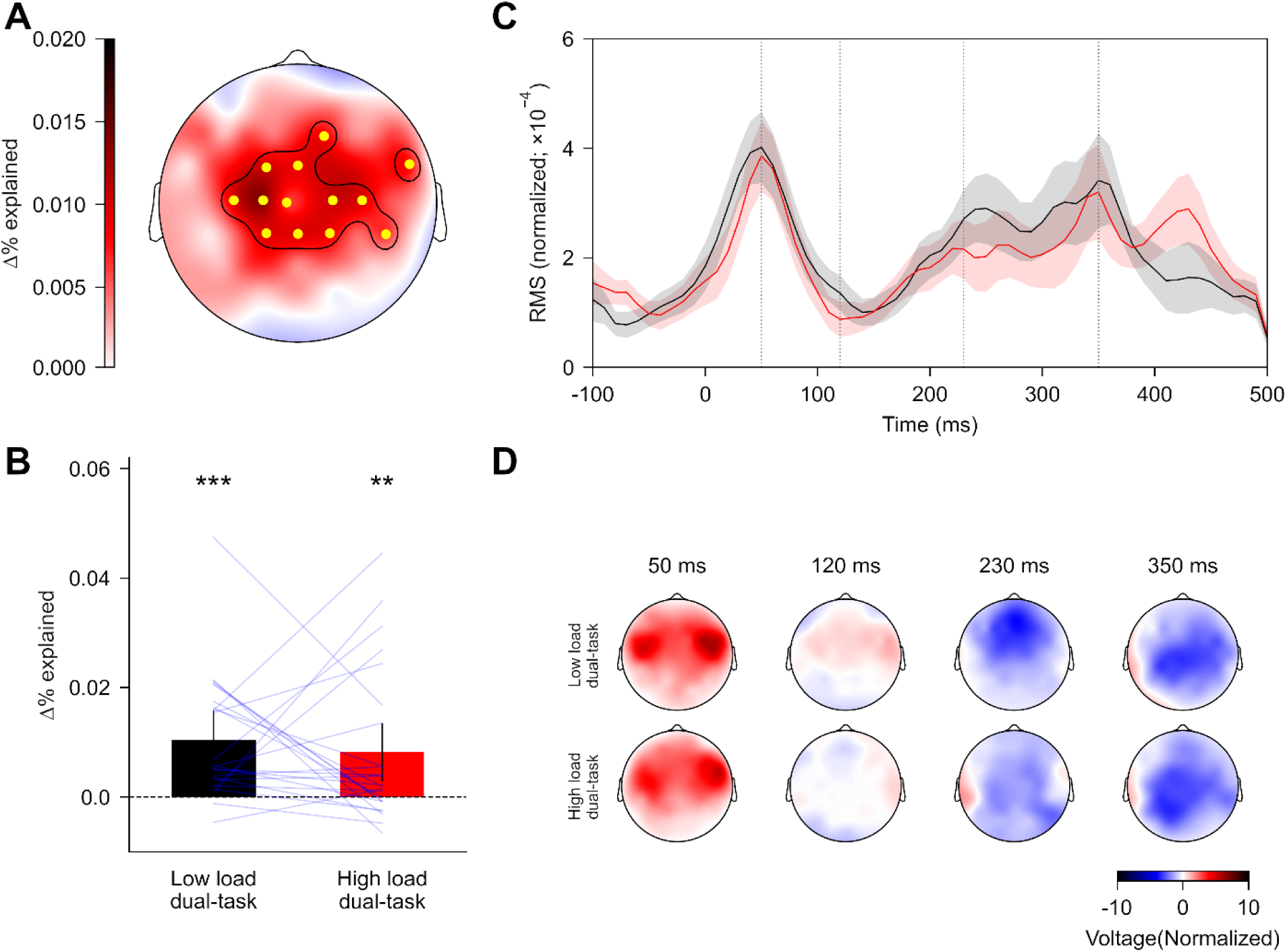
Neural tracking of word-form level linguistic predictors across dual-task conditions. (A) The topographic map illustrates the increase in prediction power (expressed as Δ% variance explained) attributed to word-form level predictors (word-form surprisal and entropy). A statistically significant increase was observed within a single cluster, marked by the highlighted yellow electrodes. (B) Prediction power across dual-task conditions, with blue lines representing individual participants. A black asterisk marks whether the prediction power for each condition was significantly above zero. Error bars reflect the 95% within-subject confidence interval. (C) The global field power of the TRFs for the word-form level predictors. The TRFs were derived by averaging responses across the surprisal and entropy predictors. Shaded regions represent within-subject standard errors around the mean (color coding as in panel B). (D) Scalp topographies at selected time points (indicated by grey vertical lines in panel C). ** *p* < 0.01, *** *p* < 0.001.

The sublexical level predictors did not significantly enhance model performance beyond visual, acoustic, and other linguistic predictors (*t*_max_ = 2.64, *p* = 0.220). These results did not provide evidence for robust neural tracking of linguistic information at the sublexical level.

In summary, the results indicate that visual task load may impair neural tracking of linguistic information at the sentence level, while leaving sublexical and word-form level processing unaffected.

### Neural tracking of acoustic features did not significantly change with visual task load

The acoustic envelope predictor significantly enhanced model performance beyond visual, acoustic onsets, and linguistic predictors, with a widespread cluster across nearly all electrodes and peak effects at temporal-central regions (*t*_max_ = 6.89, *p* < 0.001; Figure 5A). These results provide evidence for robust neural tracking of the speech acoustic envelope. Additionally, task condition did not significantly influence the prediction power for the acoustic envelope predictor [*t*(23) = 0.34, *p* = 0.365, Cohen’s d = 0.076, BF = 0.454; Figure 5B]. Finally, the mTRFs for the acoustic envelope (Figure 5C), which can be interpreted as cortical evoked responses to an elementary event in the stimulus (i.e., the impulse response), were not significantly different between the high and low load dual-task conditions.

**Figure 5.**
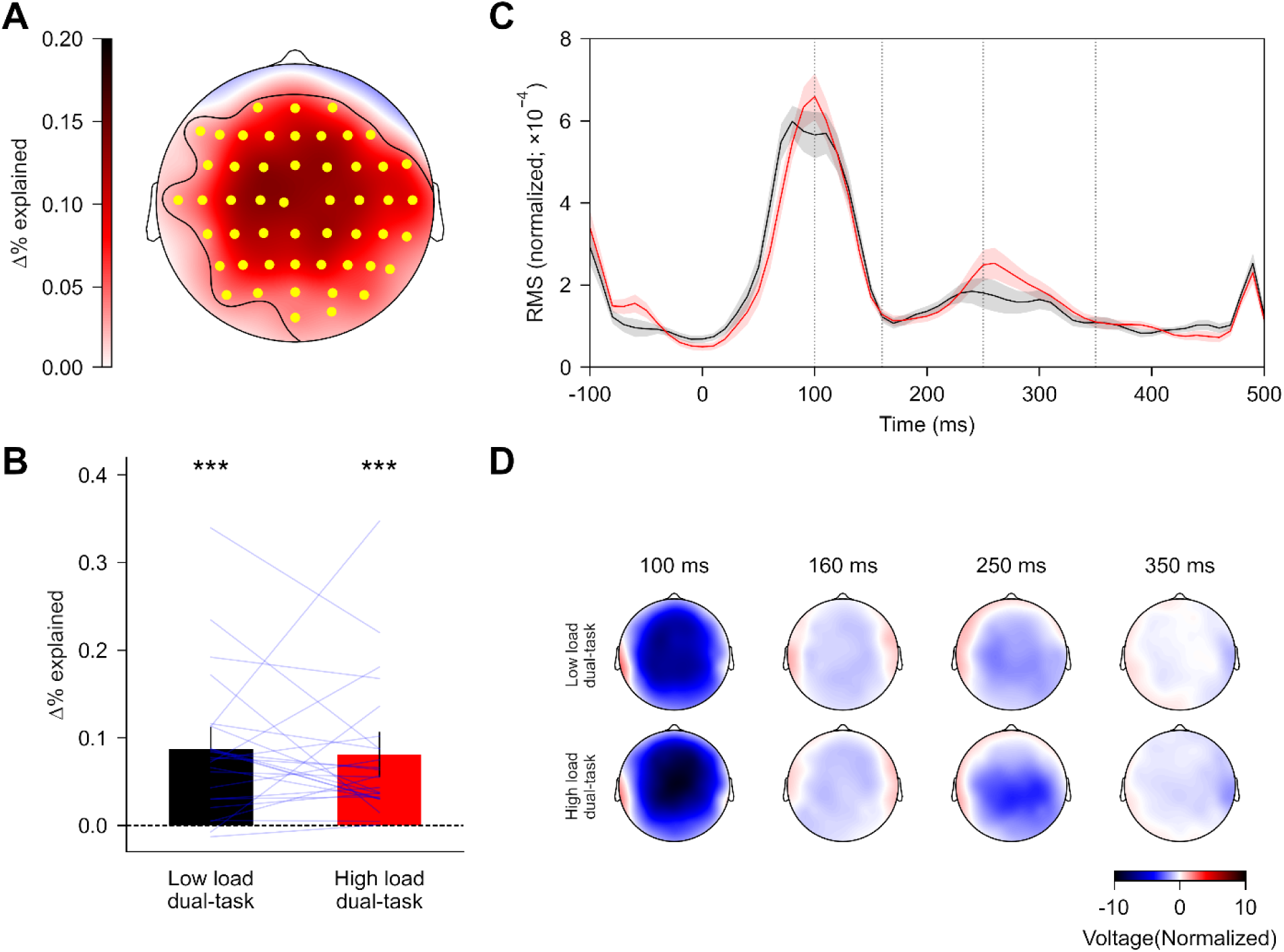
Neural tracking of the acoustic envelope of speech across dual-task conditions. (A) The topographic map illustrates the increase in prediction power (expressed as Δ% variance explained) attributed to the acoustic envelope of speech. A statistically significant increase was observed within a single cluster, marked by the highlighted yellow electrodes. (B) Prediction power across dual-task conditions, with blue lines representing individual participants. A black asterisk marks whether the prediction power for each condition was significantly above zero. Error bars reflect the 95% within-subject confidence interval. (C) The global field power of TRFs for the acoustic envelope. The TRFs were derived by averaging responses across the eight frequency bands for the envelope predictors. Shaded regions represent within-subject standard errors around the mean (color coding as in panel B). (D) Scalp topographies at selected time points (indicated by grey vertical lines in panel C). *** *p* < 0.001.

The acoustic onsets predictor significantly enhanced model performance beyond visual, acoustic envelope, and linguistic predictors, with a cluster at temporal-central regions (*t*_max_ = 6.35, *p* < 0.001; Figure 6A). These results provide evidence for robust neural tracking of the speech acoustic onsets. Additionally, task condition did not significantly influence the prediction power for the acoustic envelope predictor [*t*(23) = 1.498, *p* = 0.074, Cohen’s d = 0.332, BF = 1.147; Figure 6B]. Finally, the mTRFs for the acoustic onsets (Figure 6C) were not significantly different between the high and low load dual-task conditions.

**Figure 6.**
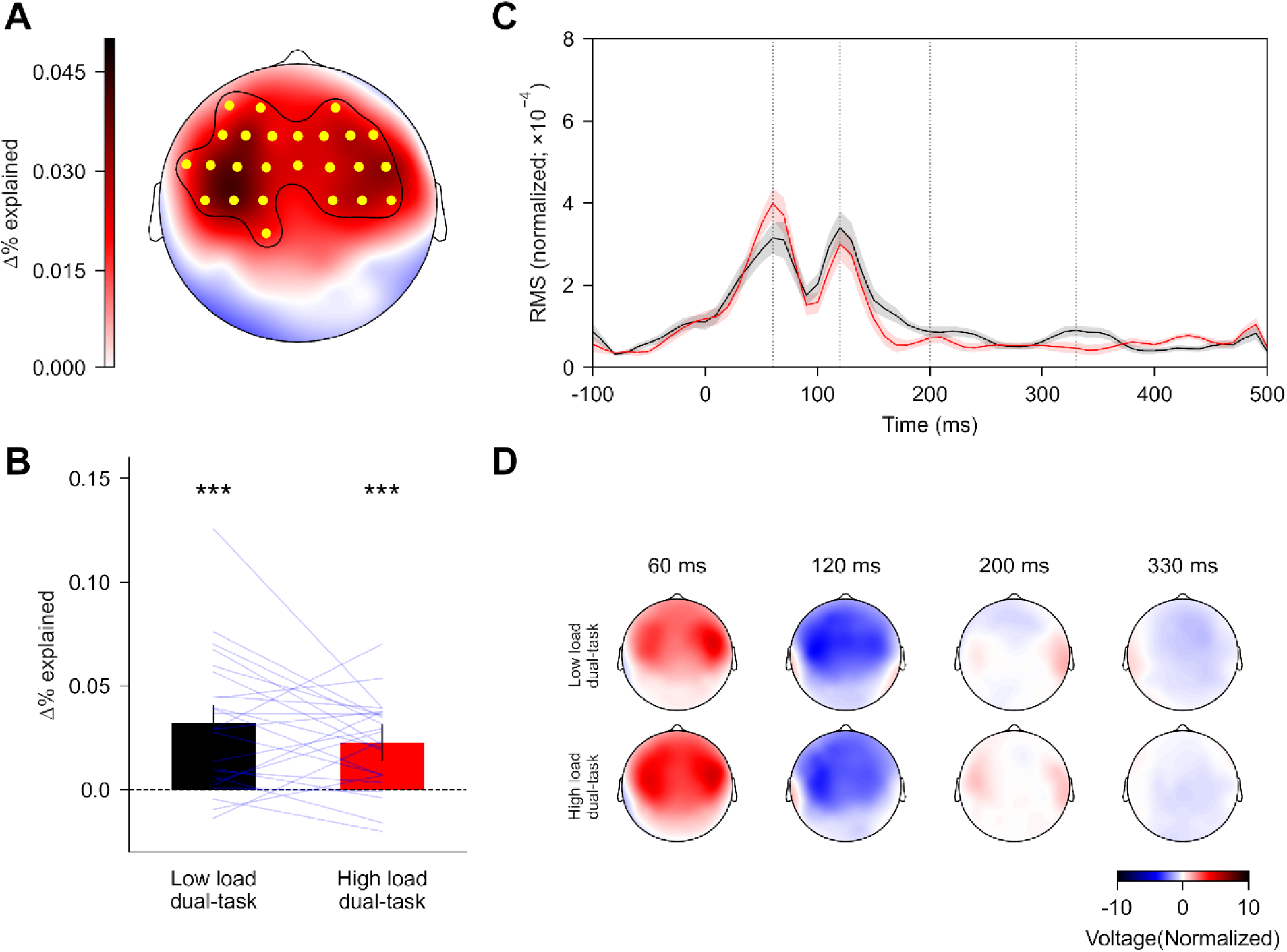
Neural tracking of the acoustic onsets of speech across dual-task conditions. (A) The topographic map illustrates the increase in prediction power (expressed as Δ% variance explained) attributed to the acoustic onsets of speech. A statistically significant increase was observed within a single cluster, marked by the highlighted yellow electrodes. (B) Prediction power across dual-task conditions, with blue lines representing individual participants. A black asterisk marks whether the prediction power for each condition was significantly above zero. Error bars reflect the 95% within-subject confidence interval. (C) The global field power of TRFs for the acoustic onsets. The TRFs were derived by averaging responses across the eight frequency bands for the onset predictors. Shaded regions represent within-subject standard errors around the mean (color coding as in panel B). (D) Scalp topographies at selected time points (indicated by grey vertical lines in panel C). *** *p* < 0.001.

To evaluate neural tracking of the visual stimuli, we examined the predictive power of visual predictors while accounting for all speech-related predictors, including acoustic and linguistic predictors. Incorporating visual predictors into a model that initially included only the speech-related predictors led to a significant increase in predictive power (*t*_max_ = 9.86, *p* < 0.001), providing evidence for neural tracking of the visual stimuli. A cluster-based analysis revealed a single significant cluster encompassing all electrodes, with the strongest effect observed in the parietal and occipital regions (Figure 7A). Furthermore, as illustrated in Figure 7B, the high load dual-task condition yielded significantly higher predictive power (mean = 0.907, *SD* = 0.504) than the low load dual-task condition (mean = 0.573, *SD* = 0.469) [*t*(23) = 3.571, *p* < 0.001, Cohen’s d = 0.684, BF = 45.993]. These results suggest that an increased visual task load enhances neural tracking of visual stimuli.

**Figure 7.**
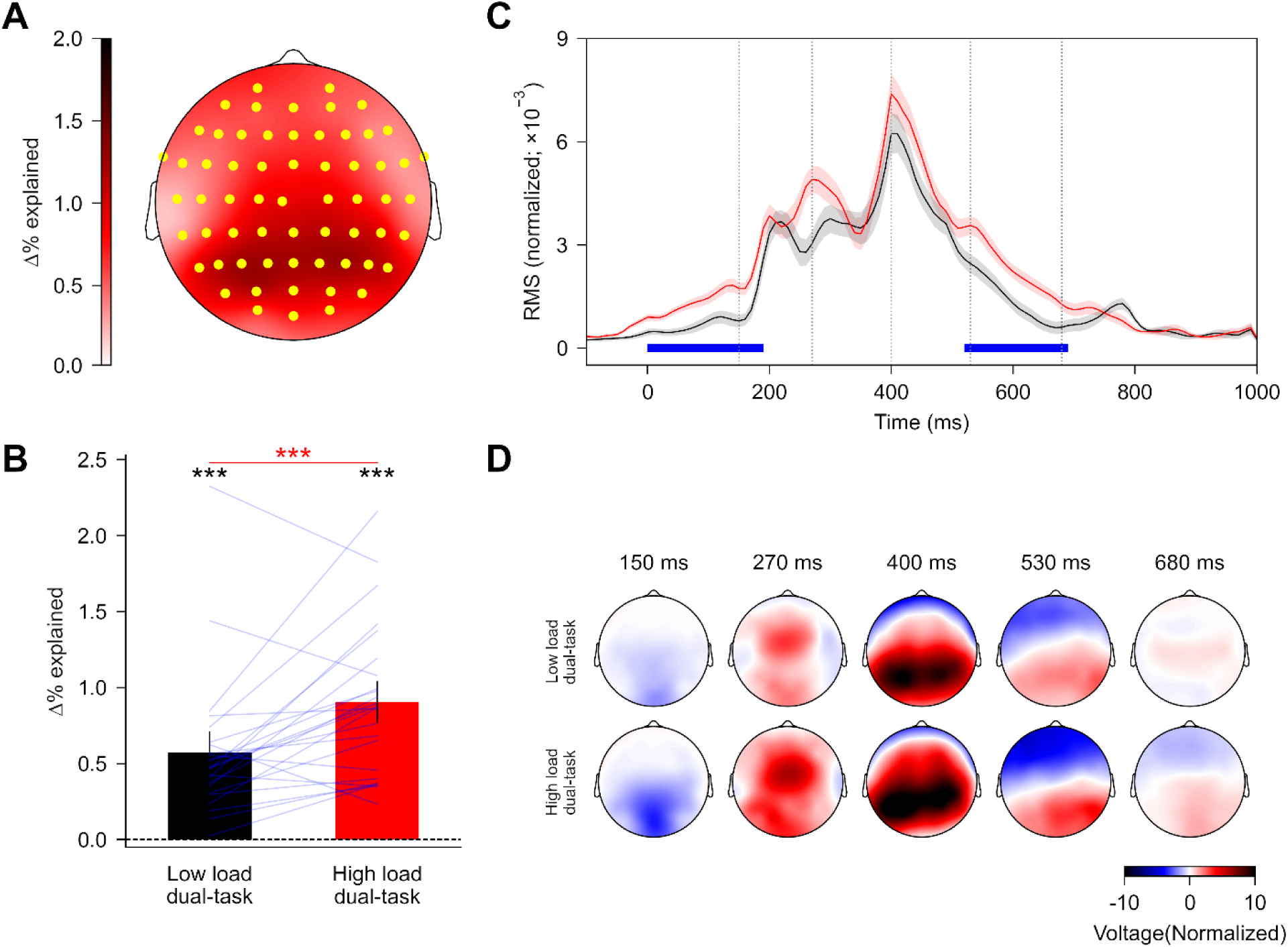
Neural tracking of visual stimuli across dual-task conditions. (A) The topographic map illustrates the increase in prediction power (expressed as Δ% variance explained) attributed to visual stimulus predictors. A statistically significant increase was observed within a single cluster, marked by the highlighted yellow electrodes. (B) Prediction power across dual-task conditions, with blue lines representing individual participants. A red asterisk indicates the statistical significance of the difference between conditions, while a black asterisk marks whether the prediction power for each condition was significantly above zero. Error bars reflect the 95% within-subject confidence interval (Loftus & Masson, 1994). (C) The global field power of the visual temporal response functions (TRFs). Each visual stimulus was displayed for a period ranging from 0 to 500 ms. Shaded regions represent within-subject standard errors around the mean (color coding as in panel B). Blue horizontal lines highlight time intervals where TRFs significantly differed between conditions. (D) Scalp topographies at selected time points (indicated by grey vertical lines in panel C). *** *p* < 0.001.

Finally, to understand how differences in model predictive power manifest in the cortical responses, we compared the mTRFs between the two dual-task conditions. These visual mTRFs can be interpreted as cortical evoked responses to the visual stimuli. In line with the findings on predictive power, as shown in Figure 7C, the high load dual-task condition produced significantly greater mTRF amplitudes than the low load dual-task condition during two time windows: from 0 to 190 ms (*p* < 0.001) and from 520 to 690 ms (*p* = 0.002).

## Discussion

The present study examined the extent to which bimodal divided attention modulates linguistic context processing at sublexical, word-form, and sentence levels during continuous speech perception in multi-talker babble. Consistent with the hypothesis, increasing dual-task load selectively reduced neural tracking of sentence-level linguistic information (Figure 3), whereas no significant effects were observed at the sublexical or word-form levels (Figure 4). Furthermore, dual-task load did not significantly alter neural tracking of acoustic features in continuous speech (Figure 5 and 6), even though both behavioral performance (Figure 2) and neural indices (Figure 7) clearly indicated that the high-load condition imposed greater processing demands.

### Hierarchical levels of linguistic context processing in quiet and noise

This study largely replicated prior findings demonstrating the parallel use of linguistic context across multiple hierarchical levels (e.g., Brodbeck et al., 2022, 2024; Karunathilake et al., 2025; Xie et al., 2023). However, unlike previous work, the current study did not observe significant cortical tracking of sublexical information. A key methodological difference is that earlier studies presented continuous speech stimuli under quiet listening conditions (e.g., Brodbeck et al., 2022, 2024; Karunathilake et al., 2025; Xie et al., 2023), whereas the present study introduced background noise. Prior research has shown that neural responses associated with sublexical context processing were relatively weaker than those observed at word-form and sentence levels (Brodbeck et al., 2022). Consequently, sublexical context processing may be more vulnerable to noise interferences, potentially accounting for the absence of significant neural tracking of sublexical information in the current study.

The central question of the present study was whether and how these multiple levels of linguistic context processing are differentially affected by bimodal divided attention. Our results showed that increasing dual-task load selectively attenuated neural tracking of sentence-level linguistic information, whereas sublexical or word-form context processing remained unaffected.

The absence of dual-task load effects at the sublexical or word-form levels is consistent with previous findings reported by Xie et al. (2023), who examined continuous speech perception under quiet listening conditions. In contrast, the selective reduction in sentence-level tracking observed here diverges from their results, which showed no dual-task load effects at this level. One possible explanation is that relative to sublexical or word-form levels, sentence-level context processing depends on the tracking and integration of linguistic information over longer timescale and therefore draws more heavily on domain-general cognitive resources (Alexander & Brown, 2018; Miller & Cohen, 2001; Shain et al., 2022). When background noise is introduced, these cognitive demands are likely amplified, rendering sentence-level processing more susceptible to the effects of increased dual-task load.

Our observation that higher-level linguistic processing is selectively reduced under increased dual-task demands appear to align with prior work using highly controlled, laboratory-based stimuli. For instance, studies examining lexical context effects on word identification within the Ganong paradigm (Ganong, 1980) have shown that increasing cognitive load via dual-task manipulations resulted in only modest changes in the lexical context effect among younger adults with normal hearing (Derawi et al., 2022; Mattys & Scharenborg, 2014). By contrast, previous research has demonstrated that neural indices of sentence-level context processing (e.g., the N400) were substantially attenuated when attention was diverted away from the language stimuli (e.g., Erlbeck et al., 2014; Hubbard & Federmeier, 2021).

### Locus of effects of bimodal divided attention on continuous speech processing

At the behavioral level, increased dual-task load clearly impaired listeners’ performance on auditory comprehension questions for continuous speech, both in noise in the current study and in quiet in Xie et al. (2023). This raises the question of where, within the speech-processing hierarchy, bimodal divided attention exerts its primary influence on behavior. Across both studies, dual-task load did not significantly modulate cortical tracking of acoustic features or lower-level linguistic representations at the sublexical or word-form level, with the only neural effects observed at the sentence level in the current study. Notably, these sentence-level effects emerged in relatively late neural responses, occurring after approximately 220 ms. Together, these findings indicate that behavioral declines under dual-task conditions predominantly reflect disruptions to later, post-perceptual stages of continuous speech processing (Xie et al., 2023).

Moreover, the sentence-level effects observed in the present study under noise conditions suggest that, when background noise is present, reduced behavioral performance may partly stem from limitations in higher-order cognitive processes involved in probabilistic word-sequence processes, such as semantic integration. Finally, the interpretation of the dual-task effects—namely, that dual-task interference in continuous speech perception primarily affects higher-order processing stages—is broadly consistent with the load theory of attention (Lavie, 2005; Lavie & Tsal, 1994). According to this framework, increases in cognitive load, such as those introduced by the visual working memory demands of our dual-task paradigm (also in Xie et al., 2023), preferentially impact later stages of stimulus processing, including memory and behavioral output (Lavie, 2005; Murphy et al., 2016; however see for example, Sörqvist et al., 2012).

### Selective versus divided attention in continuous speech processing

Prior research on auditory selective attention in continuous speech perception has shown that neural tracking of linguistic representations at sublexical or sentence levels is robust for attended speech but largely absent for unattended speech (Brodbeck et al., 2018a; Broderick et al., 2018). In contrast, findings from the present study, together with those of Xie et al. (2023), indicate that under bimodal divided attention, neural tracking at sublexical, word-form, and sentence levels is attenuated but not abolished. Notably, the effects of bimodal divided attention emerged relatively late, occurring after 200 ms. The effects of bimodal divided attention were even more limited when comparing low- and high-load dual-task conditions.

As proposed by Xie et al. (2023), these differences in attentional modulation between selective and divided attention may reflect distinct underlying mechanisms (Johnson & Zatorre, 2006; Loose et al., 2003). Selective attention costs are often attributed to “filter” mechanisms that prioritize task-relevant information while suppressing irrelevant inputs (Broadbent, 1958; Lachter et al., 2004). In contrast, the costs associated with divided attention are more likely to arise from limitations in executive control required to coordinate concurrent task demands, rather than from direct competition for shared sensory resources (Katus & Eimer, 2019; Loose et al., 2003). Nonetheless, future research is needed to more fully characterize the mechanisms that differentiate continuous speech processing under selective versus divided attention.

### Implications for domain-general cognitive processes in linguistic prediction

It remains an open question to what extent linguistic prediction during speech and language comprehension relies on domain-general cognitive processes (Ryskin et al., 2020; Ryskin & Nieuwland, 2023). The present findings, demonstrating divided attention effects on linguistic context processing in continuous speech perception, together with prior evidence from studies of selective attention, indicate that certain aspects of linguistic prediction may engage domain-general cognitive mechanisms. This appears particularly likely for predictive processes operating over longer timescales (e.g., sentence-level context) and under acoustically challenging conditions, such as in the presence of background noise. However, it remains unclear whether these cognitive processes directly modulate linguistic prediction itself or instead influence downstream measures that index predictive processing. Future research will be necessary to dissociate these possibilities and to more precisely characterize the role of domain-general cognitive processes in linguistic prediction during real-world speech comprehension.

### Study limitations

The present study employed a fixed segment of multi-talker babble as the noise and presented at a single signal-noise ratio of 5 dB. Prior research has demonstrated that the cognitive demands associated with speech perception in noise vary with both noise types (e.g., Koelewijn et al., 2012; Xie et al., 2015) and signal-to-noise ratio (e.g., Zekveld et al., 2011, 2026). Hence, it remains unclear whether the findings observed here extend to other types of background noise or to more negative or favorable signal-to-noise ratios. In addition, the extent of dual-task costs may depend on the specific characteristics of the tasks. For instance, dual-task interference may be reduced when an auditory task is paired with a visual-object task rather than a visual-spatial, such as the visuospatial n-back task used in the present study (Wahn & König, 2017). Future work should therefore examine whether the effects of bimodal divided attention on continuous speech processing vary across different combinations of auditory and visual tasks.

## Conclusion

This study suggests that bimodal divided attention may selectively disrupt cortical tracking of sentence-level linguistic representations, while leaving sublexical and word-form processing relatively unaffected during continuous speech perception in noise. Such impairments in higher-level linguistic processing may, in turn, contribute to the observed reductions in behavioral speech comprehension during multitasking in noisy environments.

## Acknowledgements

This research was funded by a startup fund from Florida State University to Z.X. The author thanks Dr. Christian Brodbeck for his valuable suggestions during the early development of the study, and Anya Chatani and Grace Frerking for their assistance with data collection and data entry. Portions of this work were presented at the 47th MidWinter Meeting of the Association for Research in Otolaryngology. The manuscript was proofread using Microsoft Copilot, after which the author conducted a thorough review and made necessary revisions, assuming full responsibility for the final content.

## Data Availability Statement

The data that support the findings of this study are available from the corresponding author upon reasonable request.

## Author Contributions

Zilong Xie: conceptualization, methodology, software, formal analysis, investigation, resources, data curation, writing–original draft, writing–review and editing, visualization, supervision, project administration, and funding acquisition.

